# Ad26-vector based COVID-19 vaccine encoding a prefusion stabilized SARS-CoV-2 Spike immunogen induces potent humoral and cellular immune responses

**DOI:** 10.1101/2020.07.30.227470

**Authors:** Rinke Bos, Lucy Rutten, Joan E.M. van der Lubbe, Mark J.G. Bakkers, Gijs Hardenberg, Frank Wegmann, David Zuijdgeest, Adriaan H. de Wilde, Annemart Koornneef, Annemiek Verwilligen, Danielle van Manen, Ted Kwaks, Ronald Vogels, Tim J. Dalebout, Sebenzile K. Myeni, Marjolein Kikkert, Eric J. Snijder, Zhenfeng Li, Dan H. Barouch, Jort Vellinga, Johannes P.M. Langedijk, Roland C. Zahn, Jerome Custers, Hanneke Schuitemaker

## Abstract

Development of effective preventative interventions against SARS-CoV-2, the etiologic agent of COVID-19 is urgently needed. The viral surface spike (S) protein of SARS-CoV-2 is a key target for prophylactic measures as it is critical for the viral replication cycle and the primary target of neutralizing antibodies. We evaluated design elements previously shown for other coronavirus S protein-based vaccines to be successful, e.g. prefusion-stabilizing substitutions and heterologous signal peptides, for selection of a S-based SARS-CoV-2 vaccine candidate. *In vitro* characterization demonstrated that the introduction of stabilizing substitutions (i.e., furin cleavage site mutations and two consecutive prolines in the hinge region of S1) increased the ratio of neutralizing versus non-neutralizing antibody binding, suggestive for a prefusion conformation of the S protein. Furthermore, the wild type signal peptide was best suited for the correct cleavage needed for a natively-folded protein. These observations translated into superior immunogenicity in mice where the Ad26 vector encoding for a membrane-bound stabilized S protein with a wild type signal peptide elicited potent neutralizing humoral immunity and cellular immunity that was polarized towards Th1 IFN-γ. This optimized Ad26 vector-based vaccine for SARS-CoV-2, termed Ad26.COV2.S, is currently being evaluated in a phase I clinical trial (ClinicalTrials.gov Identifier: NCT04436276).

## Introduction

Severe acute respiratory syndrome coronavirus 2 (SARS-CoV-2) belongs to the *Betacoronavirus* genus, the *Sarbecovirus* subgenus, and is a member of the species *SARS-related coronavirus*, and the causative agent of coronavirus disease 2019 (COVID-19) (Coronaviridae Study Group of the International Committee on Taxonomy of, 2020). In March 2020, the World Health Organization (WHO) declared the COVID-19 outbreak a pandemic, and the development of SARS-CoV-2 vaccines has become a global health priority.

Immune responses against the spike (S) protein are thought to be required for vaccine-elicited protection against coronaviruses. Like other class I fusion proteins, the SARS-CoV-2 S protein is intrinsically metastable (Sternberg and Naujokat, 2020). In recent years, efforts have been made to stabilize various class I fusion proteins in their prefusion conformation through structure-based design. In particular, the stabilization of the so-called hinge loop that precedes the central helix (CH) was shown to be a successful approach for stabilizing a range of viral fusion glycoproteins, including respiratory syncytial virus (RSV) F (Krarup et al., 2015), human immunodeficiency virus (HIV) envelope protein (Env) (Rutten et al., 2018; Sanders et al., 2002), Ebola glycoprotein (GP) (Rutten et al., 2020), human metapneumovirus (hMPV) F (Battles et al., 2017), and Lassa GP (Hastie et al., 2017). Introduction of two consecutive proline (PP) substitutions in the S2 subunit in the hinge loop between the CH and heptad repeat 1 (HR1) stabilized the S proteins of SARS-CoV and Middle East respiratory syndrome coronavirus (MERS-CoV) (Kirchdoerfer et al., 2018; Pallesen et al., 2017) and recently also the SARS-CoV-2 S protein in which also the furin cleavage site present at the boundary of the S1/S2 subunits was mutated (Walls et al., 2020; Wrapp et al., 2020). Several vaccine approaches using different designs of the S protein (e.g. with or without stabilizing substitutions, using a wild type (wt) signal peptide (SP) or Tissue Plasminogen Activator (tPA) SP (Alharbi et al., 2017)), have been described, which induce neutralizing antibodies (NAbs) and protection in animal challenge models against SARS-CoV (Martin et al., 2008), MERS-CoV (Alharbi et al., 2017) and SARS-CoV-2 infections (Corbett et al., 2020; Smith et al., 2020; van Doremalen et al., 2020; Yu et al., 2020). It is assumed that an effective vaccine against coronavirus infections should induce NAbs and a Th1-skewed immune response. For SARS-CoV prototype vaccines that were generated in response to the 2003 SARS-CoV outbreak, a theoretical risk of vaccine-associated enhanced disease induction has been associated with a Th2 type immune response in animal models (Honda-Okubo et al., 2015; Iwata-Yoshikawa et al., 2014; Tseng et al., 2012). Although it is unclear whether vaccine associated enhanced disease is a relevant concern for COVID-19 vaccines applied to humans, induction of a Th1-skewed immune response is warranted to reduce even a theoretical risk of vaccine-associated enhanced disease. Different immunogen design elements and vaccine platforms can influence these vaccine characteristics and should therefore be evaluated.

We have previously shown that vaccines based on transgenes that are delivered by recombinant replication-incompetent adenovirus type 26 vectors (Ad26) have an acceptable safety profile in humans and are able to induce neutralizing and binding antibodies, CD4 and CD8 T cell responses and a Th1-biased immune response in animals and humans (Anywaine et al., 2019; Baden et al., 2016; Baden et al., 2013; Barouch et al., 2013; Barouch et al., 2018; Liu et al., 2008; Milligan et al., 2016; Mutua et al., 2019; Radosevic et al., 2010; Shukarev et al., 2017; van der Fits et al., 2020; Winslow et al., 2017). Furthermore, the availability of industrialized and scalable manufacturing processes makes this an attractive platform for vaccine development.

Here we describe how we employ this vaccine platform for the development of an optimized S protein-based vaccine candidate against SARS-CoV-2.

## Results

### SARS-CoV-2 S immunogen design

We engineered a series of plasmids coding for seven variants of the SARS-CoV-2 S protein: 1) native full-length S (S) with wt signal peptide (SP), 2) full-length S with the tissue plasminogen activator SP (tPA.S), 3) full-length S with the tPA SP positioned upstream of the wt SP (tPA.WT.S) (Alharbi et al., 2017), 4) full-length S with the tPA SP, the furin cleavage site mutations (R682S, R685G) and proline substitutions (K986P, V987P) (tPA.S.PP)), 5) full-length S with wild type SP, with furin cleavage site mutations and proline substitutions (S.PP), 6) full-length S with only the furin cleavage site mutations (S.dFurin) and 7) full-length S with only the proline substitutions (S.PP-PR), which can therefore be processed (PR) at the furin cleavage site in the presence of furin (Fig. 1). Constructs 6 and 7 were used to study the effects of the furin cleavage site mutations and the proline substitutions separately *in vitro*.

**Figure 1.**
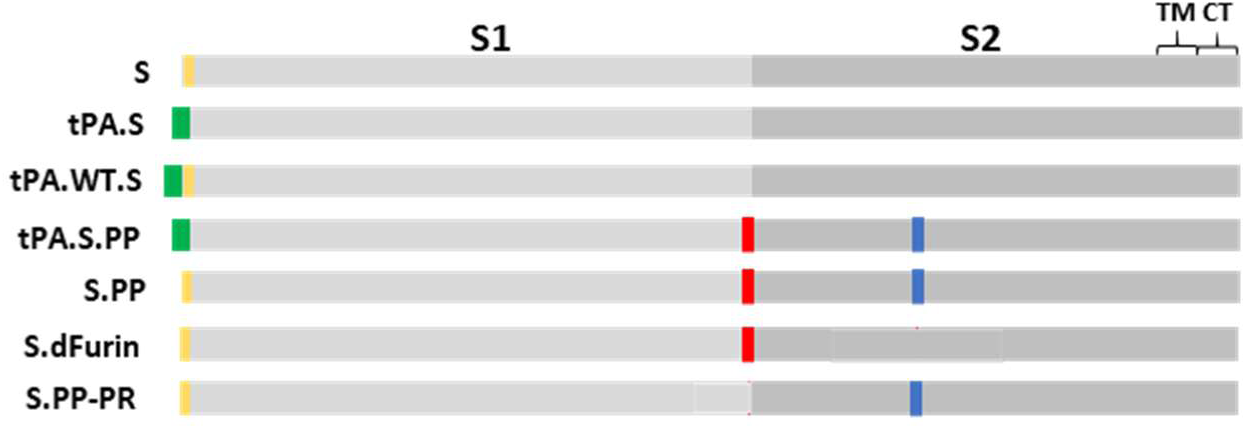
S immunogen designs. Seven plasmids encoding variants of the SARS-CoV-2 S protein were produced: 1) native full-length S (S), 2) full-length S in which the wt SP is replaced by tissue plasminogen activator SP (tPA.S), 3) full-length S in which tPA SP is added upstream of the wt SP (tPA.WT.S), 4) full-length S in which SP is replaced by tPA and in which the furin cleavage site mutations and proline substitutions (K986P, V987P) have been introduced (tPA.S.PP), 5) full-length S in which furin cleavage site mutations and proline substitutions have been introduced (S.PP), 6) full-length S with only the furin cleavage site mutations (S.dFurin) and 7) full-length S with only the proline substitutions (S.PP-PR), which is therefore processed (PR) at the furin cleavage site. Green bars represent tPA SP, yellow bars represent wild type SP, red vertical lines represent furin cleavage site mutations and blue vertical lines represent proline substitutions.

### Antigenicity of SARS-CoV-2 S designs

Antigenicity of the seven membrane bound S proteins encoded by the different DNA constructs was evaluated in cell-based ELISA (CBE). In addition, Ad26 vectors with a transgene cassette encoding five of the DNA constructs were used to transduce cells to assess transgene antigenicity by flow cytometry (Fig. 2A). Binding was assessed to four neutralizing ligands (human angiotensin-converting enzyme 2 (ACE2), convalescent human serum and two monoclonal antibodies (mAbs), S309 (Pinto et al., 2020) and SAD-S35 (Acro Biosystems) that bind the receptor binding domain (RBD)) and three mAbs (CR3022, CR3015, CR3046) that are non-neutralizing (Fig. S1). ACE2 and the neutralizing mAbs (1 μg/ml) showed the highest binding to the constructs that contain the double proline substitutions and/or the furin cleavage site mutations (tPA.S.PP, S.PP, SdFurin and S.PP-PR) (Fig. 2A). ACE2 binding to these four constructs correlated with increasing ACE2 concentration, whereas ACE2 binding to the three constructs without any stabilizing substitutions showed decreased binding at the highest ACE2 concentration (5 µg/ml) (Fig. 2B), which might be caused by shedding of the S1 domain (Tortorici and Veesler, 2019). Shedding of S1 from non-stabilized S also occurs in the absence of ACE2, as ACE2 binding activity was observed in the supernatants of cell cultures transfected with construct S, but not with S.PP (Fig. S2).

**Figure 2.**
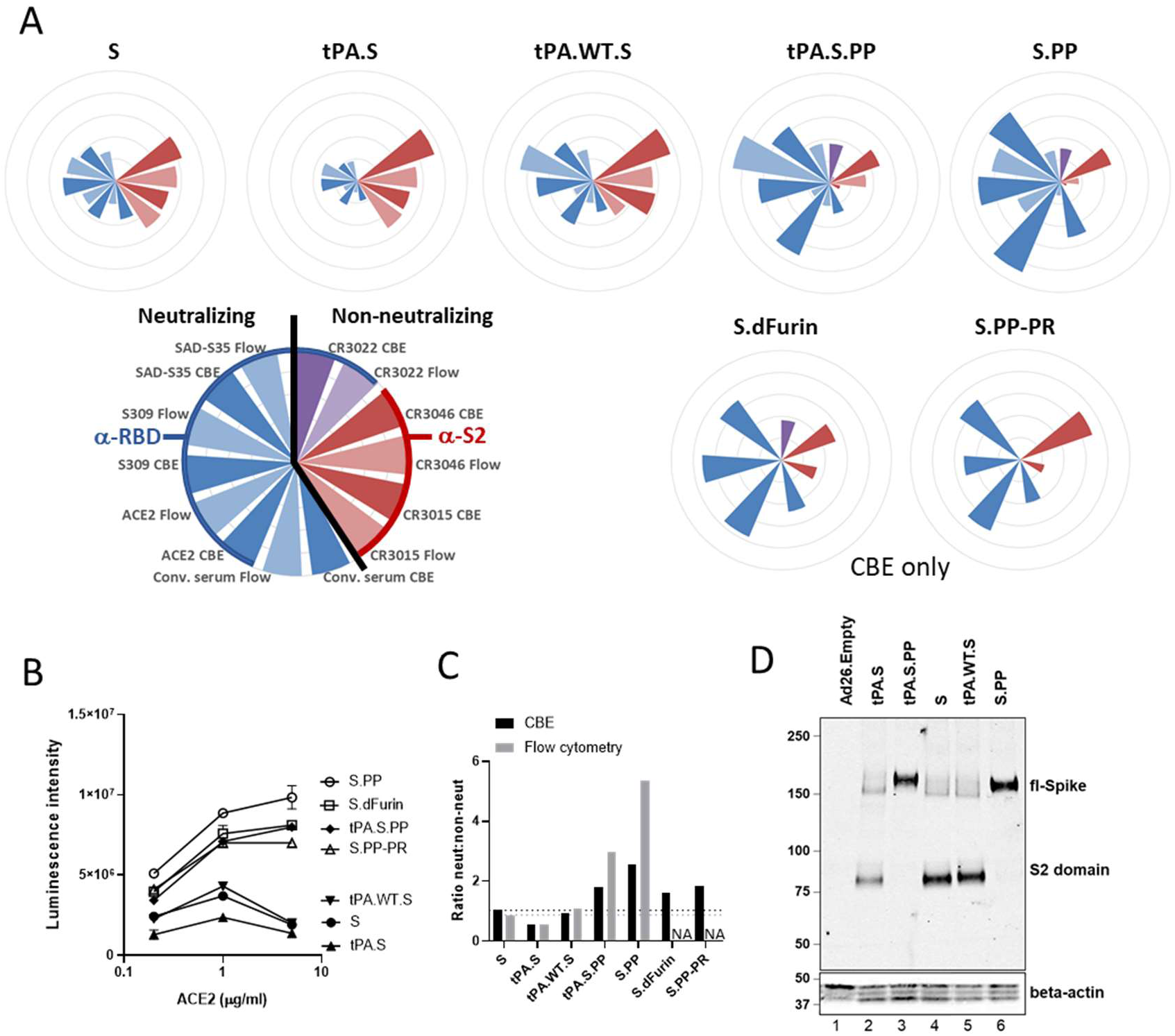
S protein antigenicity. **A**) Radar plots showing luminescence intensities measured with cell-based ELISA (CBE) and median fluorescence intensity (MFI) measured with flow cytometry (“Flow”). For CBE, cells were transfected with DNA constructs, whereas for flow cytometry cells were transduced with Ad26 vectors. Luminescence and MFI were calculated as an average of a duplicate and the MFI were normalized (by multiplying with a factor 342) to have the highest intensity the same as the highest CBE luminescence intensity. The outer ring of the circles represents a value of 10,000,000. Conv. serum stands for convalescent serum. S.dFurin and S.PP-PR were measured only with CBE. **B**) ACE2 binding measured with CBE (n=2). Data are represented as mean +SD. **C**) Average luminescence intensities measured with CBE or MFI measured with flow cytometry of neutralizing ligands and antibodies divided by that of non-neutralizing antibodies. The black horizontal dashed line indicates the height of the neut:non-neut ratio for S in CBE and the grey one for S in flow cytometry. **D**) Western blot analysis for expression from Ad26 vaccine vectors encoding tPA.S, tPA.S.PP, S, tPA.WT.S, and S.PP in MRC-5 cell lysates under non-reduced conditions using a human monoclonal antibody (CR3046). Ad26.Empty was included as negative control, and beta-actin is used as loading control

The construct containing the wt SP (S, S.PP, S.dFurin and S.PP-PR) gave a higher ratio of neutralizing to non-neutralizing antibody binding than the tPA SP (tPA.S and tPA.S.PP) (Fig. 2C). The highest ratio of neutralizing to non-neutralizing antibody binding was observed for S.PP indicating a native prefusion conformation.

Expression of all immunogens was confirmed by western blot (Fig. 2D) using the CR3046 monoclonal antibody. Furin cleavage product S2 was only seen for those constructs with an intact furin cleavage site (confirming previous observations, Mercado et al., in press).

### N-termini of mature S proteins

Correct N-terminal processing by signal peptidase is a requirement for obtaining natively-folded proteins. In coronavirus S proteins, a conserved cysteine (Cys15 in case of SARS-CoV-2 S) is present directly downstream of the SP, that forms a disulfide bond with Cys136 and which is likely required for correct folding of the N-terminal domain (NTD). *In silico* modeling using SignalP-5.0 predicted that the wt SP would be cleaved predominantly after position 15, preventing the formation of the disulfide and leading to the presence of an unpaired Cys in the mature protein. When the wt SP was replaced by the tPA SP, the predicted cleavage occurred after position 13, allowing formation of the disulfide bond to Cys136. When a tPA SP was added upstream the wt SP, SignalP-5.0 predicted cleavage after the tPA SP. To investigate the SignalP-5.0 predictions and study the effect of different SPs on S expression and stability in more detail, we performed liquid chromatography-mass spectrometry (LC-MS/MS) to determine the N-terminus of the full-length S protein as pulled down from cell membranes using either Mab CR3022 or ACE2-Fc. In contrast to the prediction of SignalP-5.0, we found that the wt SP was cleaved after position 13, that tPA SP leads to a lower number of correct N-termini and that tPA.WT.S is cleaved predominantly after position 13 of the wt SP, but that also a small fraction of incorrectly processed signal peptide was observed with the wt SP still attached (Fig. 3).

**Figure 3.**
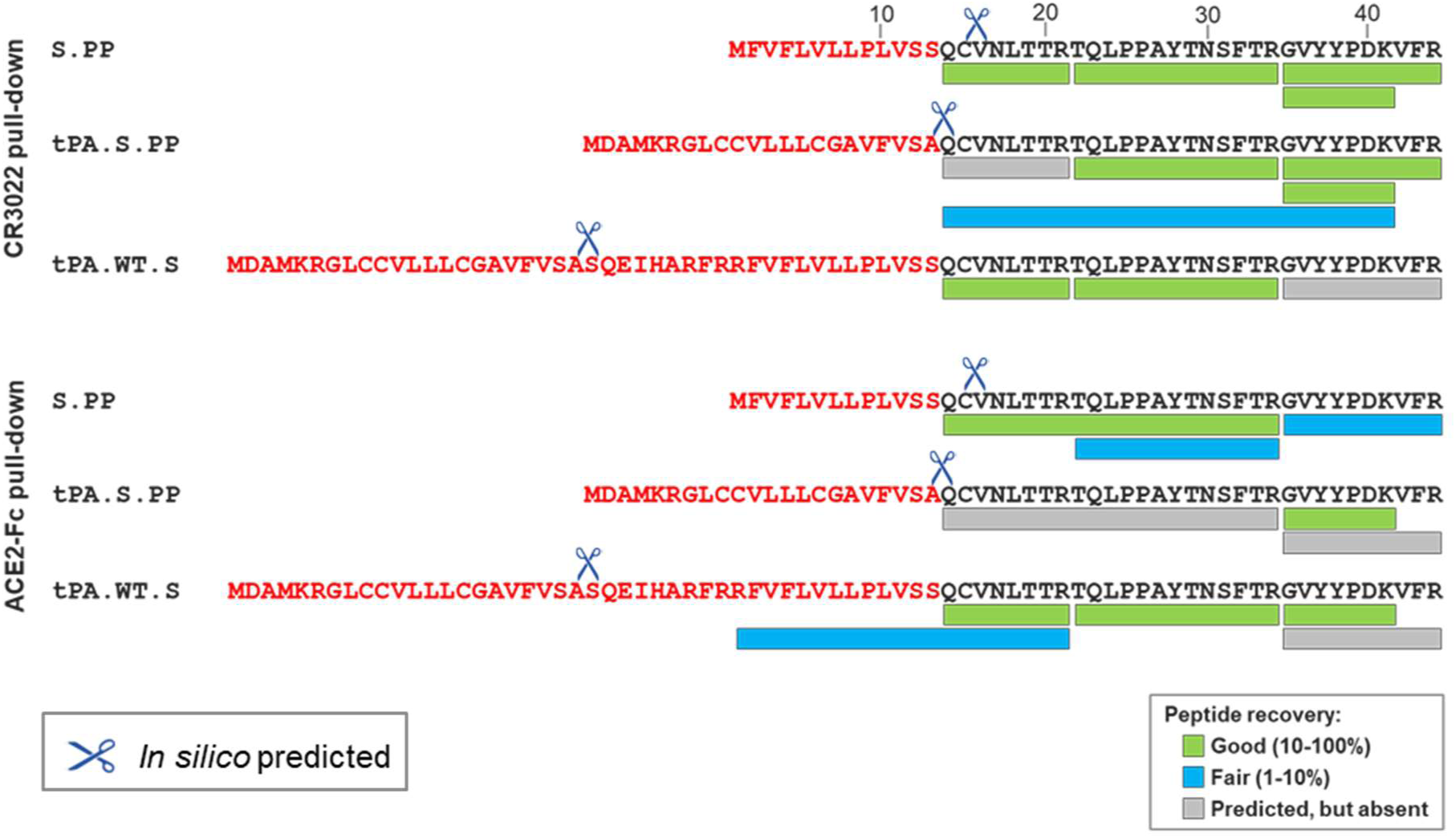
Signal peptide analysis of membrane bound S proteins. LC-MS/MS analysis of the N-terminus of S.PP, tPA.S.PP and tPA.WT.S after either ACE2-Fc or CR3022 pull-down from cell membranes. Sequences are shown of the wt SP, tPA SP and the tPA SP linked to the wt SP (in red letters) upstream of the mature S protein sequence (in black letters). Cleavage events as predicted by SignalP5.0 are indicated with scissors. The recovery rates of the different expected peptides are color-coded.

Since the membrane-bound S protein samples gave relatively low LC-MS/MS signals, probably due to inherent complexities associated with the pull-down of membrane-anchored proteins, we sought to confirm the identity of the N-terminal residue by analyzing a pair of secreted SARS-CoV-2 S proteins that differ solely in their signal peptide. To this end we purified and characterized engineered soluble proteins that were preceded by either the wt SP or the tPA SP (S.dTM.PP and tPA.S.dTM.PP, respectively). These proteins were stabilized by furin cleavage site mutations, introduction of the two consecutive proline substitutions in the S2 hinge loop and were equipped with a T4 fibritin foldon trimerization domain. Analysis of the purified soluble proteins showed that the wt SP leads to a larger fraction of correctly cleaved N-termini compared to the tPA SP (99.5% versus 11%) (Fig. S3A). In line with this, we observed lower trimer yields (Fig. S3B) and a lower melting temperature (Fig. S3C) for the protein with the tPA SP compared to the protein with the wt SP.

### Stabilizing mutations render the S protein non-fusogenic

To further characterize our immunogen designs, we determined the effect of the furin cleavage site mutations at the S1/S2 boundary as well as the PP substitutions in the S2 hinge loop on fusion competence. To this end we developed an *in vitro* cell-cell fusion assay that is based on co-transfection of constructs expressing wt SARS-CoV-2 S, and variants thereof, with constructs expressing ACE2, TMPRSS2 and green fluorescent protein (GFP) in HEK293 cells (Fig. 4A). The monolayers were imaged after 24 hr to visualize the syncytia through GFP redistribution. Both the dFurin mutations and the PP substitutions alone or in combination, were sufficient to fully prevent syncytium formation (Fig. 4B).

**Figure 4.**
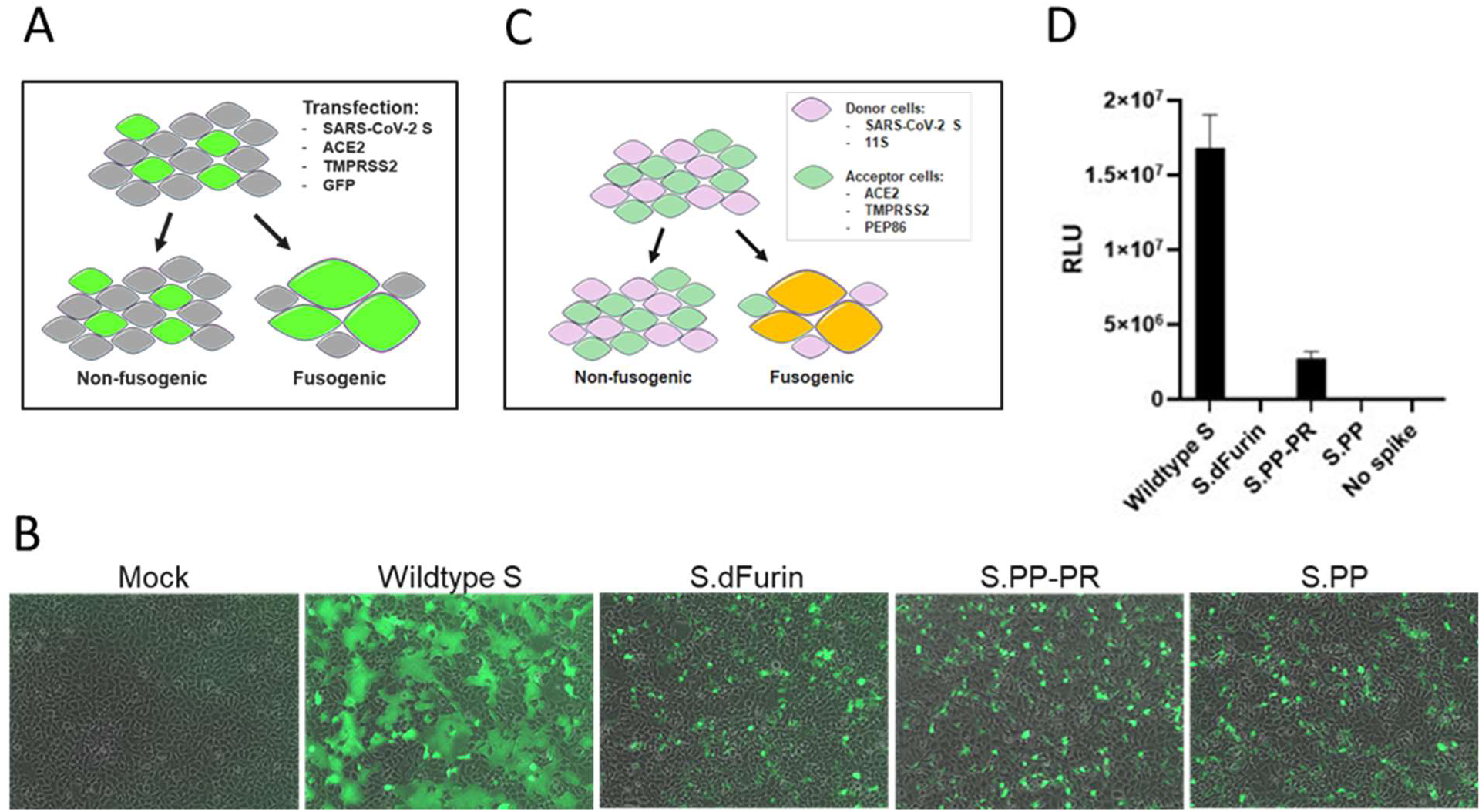
Effect of stabilizing mutations on fusogenicity of S protein. **A**) S protein fusogenicity as measured in a cell-cell fusion assay in HEK293 cells by co-transfection of plasmids encoding S protein, ACE2, TMPRSS2 and GFP. **B**) Overlay of GFP and brightfield channels 24 h after transfection, as in the setup of (A). The different S protein constructs are indicated; mock is an untransfected monolayer. **C**) Quantitative cell-cell fusion assay setup. **D**) Luciferase signal shown as relative light units (RLU) measured at 4 h post mixing of donor and acceptor cells, as in the setup of (C).

Next, to increase sensitivity and allow quantification of the fusion assay, we redesigned the assay using a split luciferase system. A donor cell line, expressing the S protein and one luciferase subunit, was mixed with an acceptor cell line that expresses ACE2, TMPRSS2 and the complementing luciferase subunit (Fig. 4C). In this setup, expression of the wt S protein yielded high signals that were diminished upon introduction of the PP substitutions but remained detectable (Fig. 4D). Furin cleavage site mutations reduced the signal to the background level (‘no spike’). As expected, combination of both stabilizing mutations rendered the S protein non-fusogenic as well.

### Stabilizing substitutions enhance in vivo immunogenicity

In parallel to the *in vitro* characterization, we assessed how the different construct design elements affect immunogenicity. Mice received a single intramuscular (i.m.) immunization of 10^8^, 10^9^ or 10^10^ viral particles (vp) of Ad26 vectors encoding different variants of the SARS-CoV-2 S protein. We observed that across doses, adeno26 with the tPA.S insert (Ad26.tPA.S) induced lower titers of antibodies that bind to the S protein than Ad26.S, as measured by ELISA at 4 and 6 weeks post immunization (Fig. 5A,B). Ad26.tPA.S.PP and Ad26.tPA.WT.S showed similar immunogenicity as Ad26.S. In addition, a live virus neutralization assay (VNA) was used to measure SARS-CoV-2 NAb responses in mice receiving 10^10^ vp (Ad26.tPA.S was not included). Ad26.tPA.S.PP did not induce higher NAb titers than Ad26.S (Fig. 5C,D), in line with the findings described above. Ad26.tPA.WT.S induced lower NAb titers than Ad26.S albeit not statistically significant.

**Figure 5.**
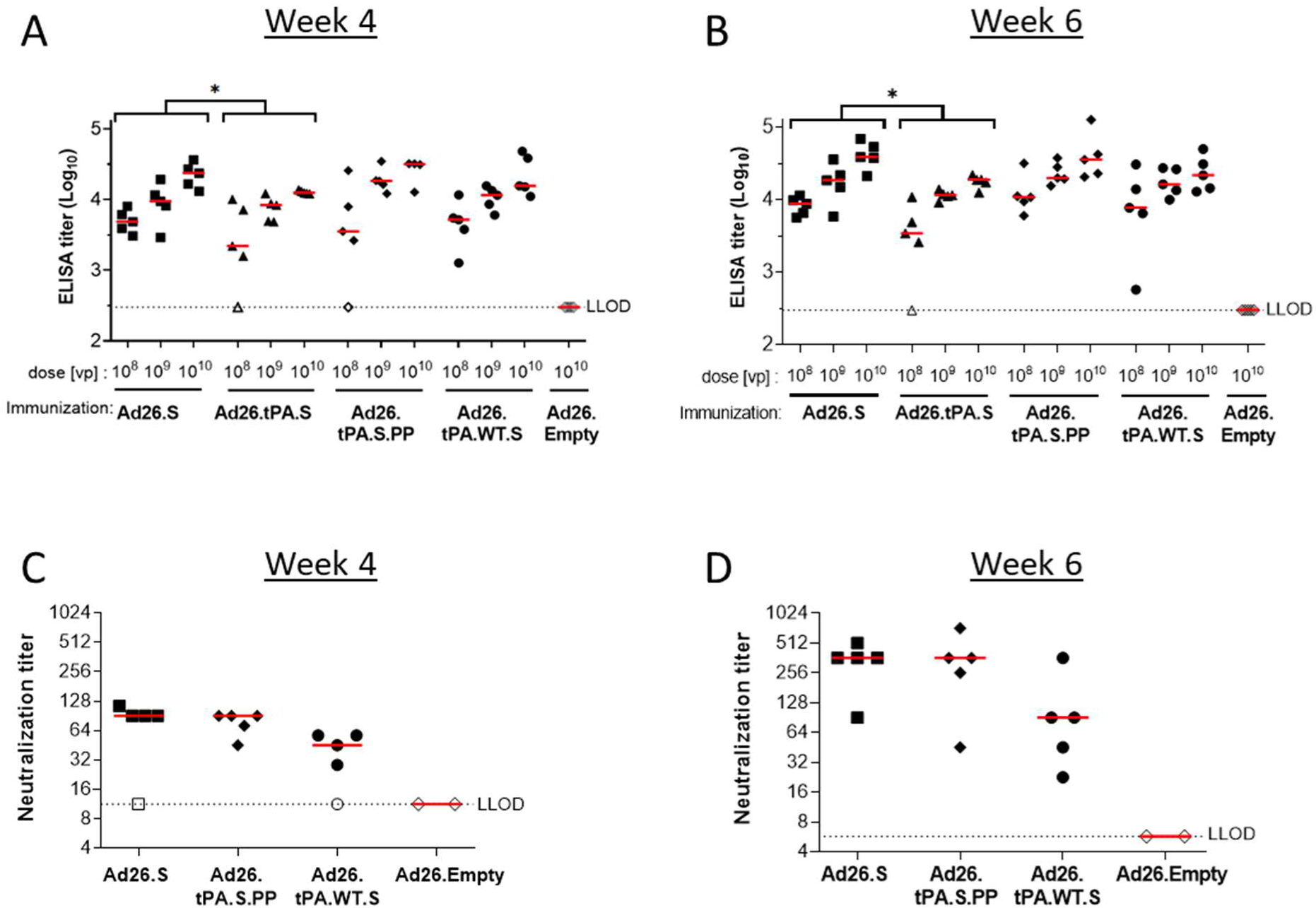
Immunization with Ad26-based vaccine constructs induce humoral immune responses in immunized mice. Naïve mice (C57BL/6, N=5 per group) were immunized with either 10^8^, 10^9^ or 10^10^ vp of Ad26-based vaccine candidates, or with 10^10^ vp of an Ad26 vector without gene insert as control (Ad26.Empty). Four weeks and six weeks after the immunization, S protein-specific binding and NAb titers were determined. **A + B**) Spike protein-specific antibody binding titers were measured by ELISA. **C + D**) SARS-CoV-2 NAb titers were measured by wt VNA determining the inhibition of the cytopathic effect (CPE) of virus isolate Leiden1 (L-0001) on Vero E6 cells. Mice immunized with Ad26.empty were taken along as two separate pools. Median responses per group are indicated with horizontal lines. Dotted lines indicate the LLOD per assay. Animals with a response at or below the LLOD were put on LLOD and are shown as open symbols. Statistical differences are indicated by asterisks; *: p<0.05. LLOD = Lower Limit of Detection. vp = virus Particles.

We next compared humoral and cellular immune responses induced by Ad26.S.PP and Ad26.S. Mice received a single i.m. immunization of 10^8^, 10^9^ or 10^10^ vp Ad26.S or Ad26.S.PP and binding and NAbs titers were measured at weeks 2 and 4. Both vectors elicited S protein binding antibodies in a dose-dependent manner (Fig. 6A,B). The induction of Nabs was only assessed for mice immunized with the highest dose (Fig. 6 C,D). Titers of both binding- and NAbs were significantly higher in Ad26.S.PP immunized mice, clearly indicating the benefit of S protein stabilization for immunogenicity. These findings were consistent with immunizations with DNA vaccines encoding tPA.S, tPA.S.PP, S and S.PP (Fig. S4).

**Figure 6.**
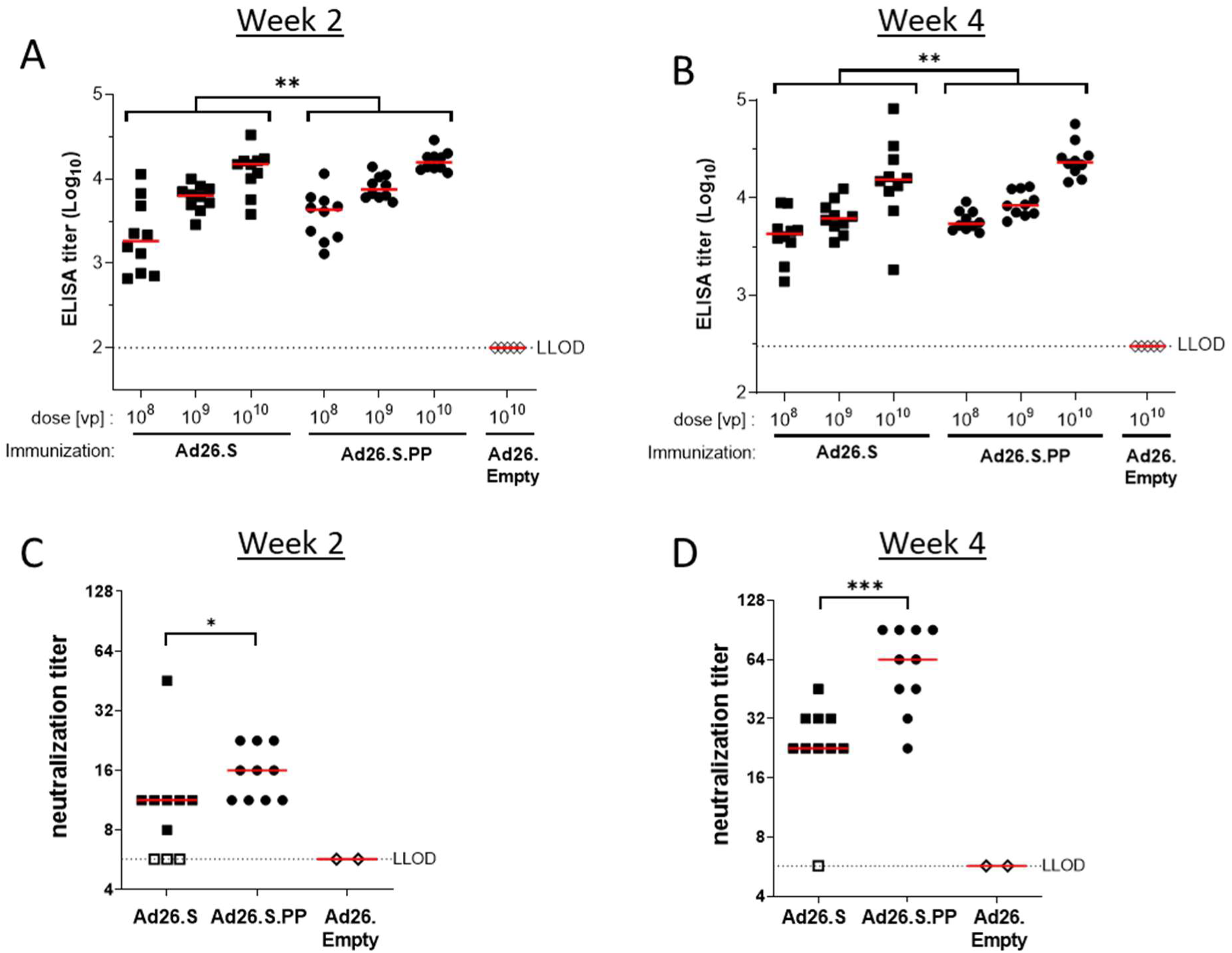
Significant higher S protein binding antibody titers and SARS-CoV-2 neutralization titers induced by Ad26.S.PP compared to Ad26.S. Naïve mice (BALB/c, N=10 per group) were immunized with either 10^8^, 10^9^ or 10^10^ vp of Ad26.S.PP, Ad26.S or with Ad26.Empty (N=5). Serum was sampled 2 and 4 weeks post immunization and splenocytes were analyzed 4 weeks post immunization. **A + B)** S protein-specific binding antibody titers were measured by ELISA. **C + D)** SARS-CoV-2 NAb titers were measured by wt VNA. Mice immunized with Ad26.empty were taken along as two separate pools in VNA. Median responses per group are indicated with horizontal lines. Dotted lines indicate the LLOD per assay. Animals with a response at or below the LLOD were put on LLOD and are shown as open symbols. Statistically significant differences between groups per dose, or across doses (indicated by brackets), are indicated by asterisks; *: p<0.05, **: p<0.01; ***p<0.001. LLOD = Lower Limit of Detection. vp = virus Particles.

### Ad26.S.PP induces Th1-skewing of T cell response

To further investigate whether Ad26.S.PP induced a Th1-skewed immune response, to reduce the theoretical risk of vaccine-associated enhanced disease, mice were immunized i.m. with a single dose of 10^8^ or 10^10^ vp Ad26.S or Ad26.S.PP and T cell responses were measured by IFN-γ ELISpot after 4 weeks. Both vaccines elicited IFN-γ producing T cells and no differences were observed between doses or vaccine constructs (Fig. 7A). To further assess the Th1/Th2 polarization by our lead vaccine candidate, mice were immunized with either Ad26.S.PP or 50 µg recombinant S protein in 100 µg aluminum phosphate adjuvant (Adjuphos), which has been shown to induce a Th2-biased immune response (Lindblad, 2004). A single dose of Ad26.S.PP again elicited IFN-γ producing T cells as measured by ELISpot. However, recombinant S protein in Adjuphos induced undetectable or low IFN-γ ELISpot responses (Fig. 7B). Th1 dominant T cell responses were confirmed by ICS (Fig. S5). Stimulation of splenocytes with S peptide pools resulted in secretion of the Th1 hallmark cytokine IFN-γ, but secretion of typical cytokines associated with a Th2-type immune response was low. The ratios of IFN-γ concentration to either IL-4, IL-5, or IL-10 concentration was high after immunization with Ad26.S.PP suggesting that the response was Th1-skewed (Fig. 7C). IgG1 is produced during any type of immune response, while IgG2a is predominantly produced during a Th1-polarized immune response, and therefore an increase of the IgG2a/IgG1 ratio is indicative of the Th1-skewing of the immune response. Ad26.S.PP induced a high IgG2a/IgG1 ratio again indicative of a Th1-biased response, whereas S protein in Adjuphos induced a low IgG2a/IgG1 ratio, confirming a Th2-biased response (Fig. 7D).

**Figure 7.**
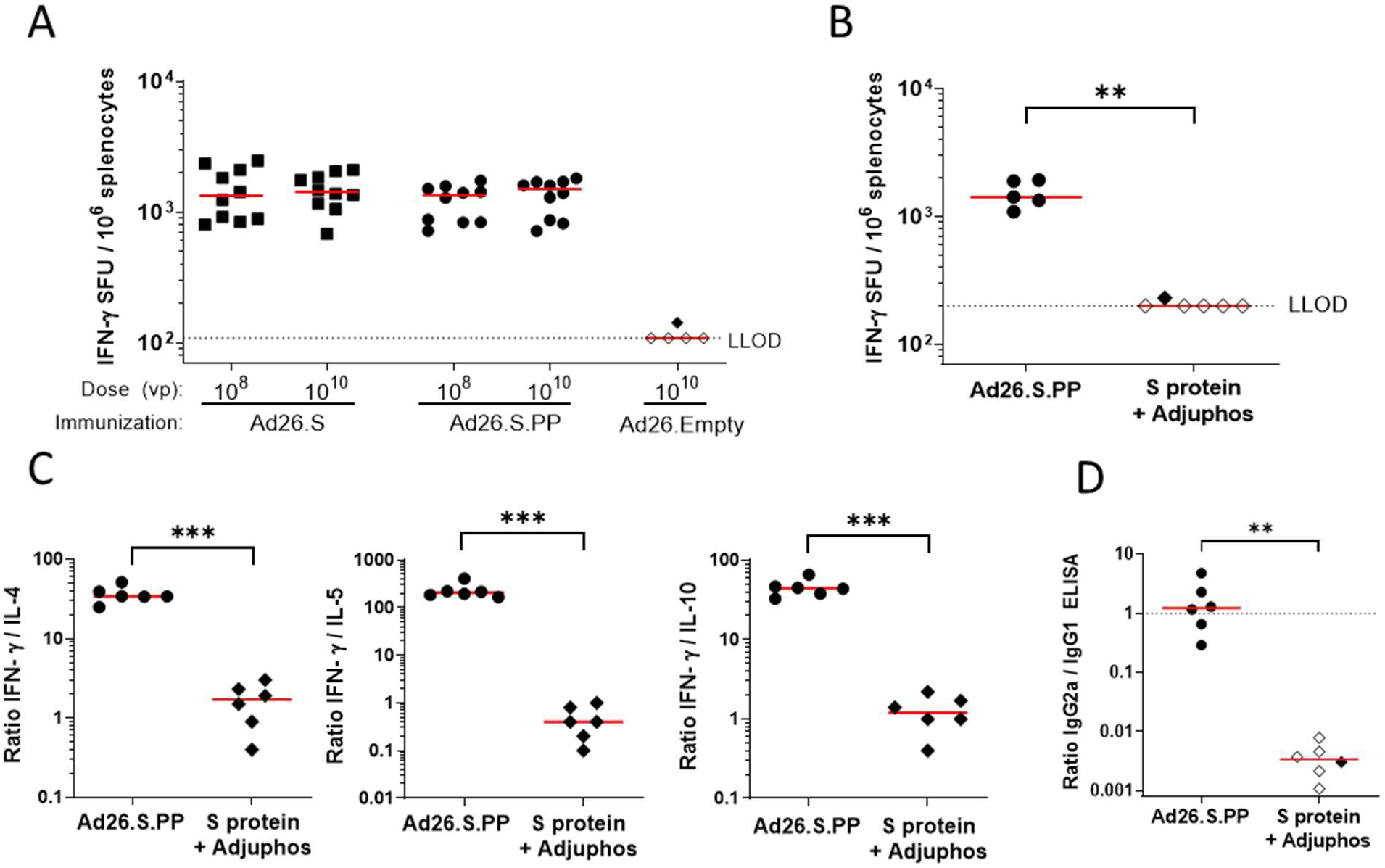
Analysis of IFN-γ secretion and Th1/Th2 skewing in Ad26.S.PP immunized mice. **A)** Naïve BALB/c mice were immunized with either 10^8^ or 10^10^ vp of Ad26.S.PP, Ad26.S (N=10 per group) or with Ad26.Empty (N=5). Number of IFN-γ producing cells per million splenocytes four weeks after immunization as measured by ELISpot. Median responses per group are indicated with horizontal lines. Dotted line indicates the LLOD. Animals with a response at or below the LLOD were put on LLOD and are shown as open symbols. **B, C, and D)** Naïve mice (BALB/c, N=6 per group) were immunized with either 10^10^ vp of Ad26.S.PP or 50 µg of S protein adjuvanted with 100 µg Adjuphos (Adju-Phos^®^). Two weeks after immunization samples were analyzed for antibody and cellular responses. **B)** Number of IFN-γ producing cells per million splenocytes as measured by ELISpot. One mouse excluded due to high background in peptide pool 1 stimulation. Dotted line indicates the LLOD. **C)** Th1 (IFN-γ) over Th2 (IL-4, IL-5 and IL-10) cytokine ratios measured by multiplex ELISA after stimulation of splenocytes with SARS-CoV-2 S protein peptides. Ratio of IgG2a/IgG1 antibody titers as measured by IgG2a and IgG1 ELISA. Dotted line indicates an IgG2a/IgG1 ratio of 1. **D)** Ratio of IgG2a/IgG1 antibody titers as measured by IgG2a and IgG1 ELISA. Dotted line indicates an IgG2a/IgG1 ratio of 1. Animals with a response at or below the LLOD in the IgG2a ELISA are shown as open symbols. Horizontal lines denote group medians. Statistically significant differences indicated by asterisks **: p<0.01; ***p<0.001. LLOD= Lower Limit of Detection. vp= virus Particles.

## Discussion

According to the WHO, more than 200 COVID-19 vaccine candidates are currently in development of which most target SARS-CoV-2 S protein. Our data demonstrate that the design of the S protein is critical for the characteristics of the immunogen and the resulting immune responses elicited by the vaccine. While SARS-CoV-2-binding antibodies and NAbs have been shown to be induced in animal models and humans by several vaccines using different vaccine platforms and different S protein designs (Corbett et al., 2020), we here examined different S protein designs side by side which revealed specific characteristics for each of them. We show here that the wt SP is best suited for correct cleavage, which is needed for natively folded protein and it enhanced the induction of S binding antibodies and virus NAbs *in vivo*. Initial in silico analysis revealed that cleavage after the wt SP could result in the removal of the structurally important cysteine at position 2 of the mature protein. However, replacement of the wt SP by the tPA SP resulted in aberrant cleavage after the conserved N-proximal cysteine, most probably required for fully correct S protein folding whereas only the wt SP resulted in fully native amino terminal cleavage. This agreed well with reduced neutralizing antibody binding to tPA.S versus S. A high ratio of neutralizing to non-neutralizing antibody binding is a qualitative measure of high expression levels of S in the prefusion conformation and/or low amount of S1 shedding (Tortorici and Veesler, 2019). The wt SP in combination with modifications that stabilize the protein in the prefusion conformation, i.e. furin cleavage site mutations and PP substitutions (S.PP), resulted in the highest ratio of neutralizing to non-neutralizing antibody binding. *In vivo* evaluation of the Ad26-based vaccines encoding the different S protein designs confirms that they all are immunogenic in mice after a single immunization. Induction of binding antibodies was observed for all Ad26 doses tested and S.PP was most immunogenic as it elicited the highest binding- and neutralizing antibody titers. This is likely related to the stabilizing effect of the PP substitutions and mutations in the furin cleavage site that preserve the prefusion conformation and blocks shedding of S1 (Fig. 2D). Furthermore, the combination of stabilizing mutations renders the S protein non-fusogenic, indicating that the conformational change to the post-fusion form is prevented, in line with improved antigenicity.

An important aspect in the development of a SARS-CoV-2 vaccine is to exclude or minimize the theoretical risk of vaccine-associated enhanced disease that was observed in animal models for some of the SARS-CoV (Honda-Okubo et al., 2015; Iwata-Yoshikawa et al., 2014; Tseng et al., 2012). In these models, vaccine-associated disease enhancement was associated with low neutralizing antibody titers and a Th2 skewed immune response. These findings, however, could not be substantiated in humans as no SARS-CoV vaccine was tested for efficacy. Genetic vaccines like Ad26 have been shown to induce a Th1-biased immune response(Anywaine et al., 2019; Baden et al., 2016; Baden et al., 2013; Barouch et al., 2013; Barouch et al., 2018; Liu et al., 2008; Milligan et al., 2016; Mutua et al., 2019; Radosevic et al., 2010; Shukarev et al., 2017; van der Fits et al., 2020; Winslow et al., 2017), and we have shown here that Ad26.S.PP elicited a dominant Th1 response in combination with high titers of Nab, thereby reducing the theoretical risk of vaccine-associated enhanced disease. Ongoing animal studies are aimed to further substantiate these findings.

The potency of the Ad26-based vaccine encoding S.PP (Ad26.S.PP) in eliciting protective immunity against SARS-CoV-2 infection was successfully demonstrated in a non-human primate challenge model (Chandrashekar et al., 2020) (Mercado et al., in press). Ad26.S.PP, from now on named Ad26.COV2.S, was identified as our lead SARS-CoV-2 vaccine candidate and is currently being evaluated in a phase I clinical trial. (ClinicalTrials.gov Identifier: NCT04436276).

## Acknowledgements

This project was funded in part by the Department of Health and Human Services Biomedical Advanced Research and Development Authority (BARDA) under contract HHS0100201700018C.

We thank Isabel de los Rios Oakes, Esmeralda van der Helm, Dirk Spek, Iris Swart, Marina Koning, Alies Brandjes, Nelie van Dijk, Eleni Kourkouta, Santusha Karia, Marjon Navis, Rina van Schie, Janneke Verhagen, Jessica Vreugdenhil, Sanne Kroos, Ella van Huizen, Jeroen Tolboom, Leacky Muchene, Joke Drijver, Liz van Erp, Michel Mulders, Aric van Drie, Eveline Sneekes-Vriese, Sharitee Bouthisma, Ava Sadi, Sven Blokland, Richard Voorzaat, Lam Le, Remko van der Vlugt, Theo Schouten, Lisa Tostanoski, Noe Mercado, and Katherine McMahan for generous advice and assistance.

## Author Contributions

Designed studies and reviewed data: R.B., L.R., J.E.M.vd.L., M.J.G.B., G.H., F.W., D.Z., R.V., A.H.d.W., M.K., E.J.S., S.K,M., J.V., J.P.M.L., R.Z., J.C., A.V., A.K., D.v.M., J.C., T.K., H.S.. Design of vaccines: L.R, M.J.G.B, R.B, A.H.d.W, D.Z, T.K, J.P.M.L, F.W, D.v.M, Z.L, R.C.Z, J.V, J.C., R.V, D.H.B., H.S.. Performed experiments: T.J.D., S.K.M. and A.K.. Drafted the paper: all authors.

## Methods

### Vaccine design

The S protein of SARS-CoV-2 corresponding to positions 21,536 to 25,384 in SARS-CoV-2 isolate Wuhan-Hu-1 (GenBank accession number: MN908947) was codon-optimized for expression in human cell lines. S designs were either based on the native SP, replacement of the native SP by the tissue plasminogen activator (tPA) SP or a SP upstream of the native signal, resulting in a sequence encoding SARS-CoV-2 amino acids 2-1273 and tPA leader as described for MERS-CoV spike protein (Alharbi et al., 2017; Gilbert et al., 2017) In some constructs the furin cleavage site was abolished by amino acid changes R682S and R685G, or proline substitutions (K986P, V987P) were introduced.

### Protein expression and purification

A plasmid encoding the SARS-CoV2 S-2P protein (Wrapp et al., 2020) with the wt SP and with the wt SP replaced by the tPA SP and with a C-tag for purification were codon-optimized and synthesized at GenScript. The constructs were cloned into pCDNA2004. The expression platform used was the Expi293F cells. The cells were transiently transfected using ExpiFectamine (Life Technologies) according to the manufacturer’s instructions and cultured for 6 days at 37°C and 10% CO_2_. The culture supernatant was harvested and spun for 5 min at 300 g to remove cells and cellular debris. The supernatant was subsequently sterile filtered using a 0.22 µm vacuum filter and stored at 4°C until use. The C-tagged SARS-CoV2 S trimers were purified using a two-step purification protocol by 5 mL CaptureSelect™ C-tag Affinity Matrix (ThermoFisher Scientific). Proteins were further purified by size-exclusion chromatography using a HiLoad Superdex 200 16/600column (GE Healthcare).

### Antibodies and reagents

SAD-S35 was purchased at Acro Biosystems. ACE2-Fc (ACE2) was made according to Liu et al. 2018 (Liu et al., 2018). For CR3022, CR3015 (van den Brink et al., 2005), CR3046 and S309 (Pinto et al., 2020) the heavy and light chain were cloned into a single IgG1 expression vector to express a fully human IgG1 antibody. The antibodies were made by transfecting the IgG1 expression construct using the ExpiFectamine™ 293 Transfection Kit (ThermoFisher) in Expi293F (ThermoFisher) cells according to the manufacturer specifications. Antibodies were purified from serum-free culture supernatants using mAb Select SuRe resin (GE Healthcare) followed by rapid desalting using a HiPrep 26/10 Desalting column (GE Healthcare). The final formulation buffer was 20 mM NaAc, 75 mM NaCl, 5% Sucrose pH 5.5. Convalescent serum (SER-0743-00001) was obtained from Sanquin, the Netherlands.

### Cell-Based ELISA

HEK293 cells were seeded at 2 × 10^5^ cells/mL in appropriate medium in a flat-bottomed 96-well microtiter plate (Corning). The plate was incubated overnight at 37 °C in 10% CO_2_. After 24 hrs, transfection of the cells was performed with 300 ng DNA for each well and the plate was incubated for 48 hrs at 37 °C in 5% CO_2_. Two days post transfection, cells were washed with 100 μl/well of blocking buffer containing 1% (w/v) BSA (Sigma), 1 mM MgCl_2_, 1.8 mM CaCl_2_ and 5 mM Tris pH 8.0 in 1x PBS (GIBCO). After washing, nonspecific binding was blocked, using 100 μl/well of blocking solution for 20 min at 4 °C. Subsequently, cells were incubated in 50 μl/well blocking buffer containing primary antibodies ACE2-Fc (5 µg/mL, 1 µg/mL and 0.2 µg/mL)(1 µg/mL for radar plot), S309 (1 µg/mL), SAD-S35 (1 µg/mL), CR3015 (5 µg/mL), CR3022 (5 µg/mL), CR3046 (5 µg/mL), and convalescent serum (1:400) for 1 hr at 4 °C. The plate was washed three times with 100 μl/well of the blocking buffer, three times with 100 μl/well of washing buffer containing 1 mM MgCl_2_, 1.8 mM CaCl_2_ in 1x PBS and then incubated with 100 μl/well of the blocking buffer for 5 min at 4 °C. After blocking, the cells were incubated with 50 μl/well of secondary antibodies HRP conjugated Mouse Anti Human IgG (Jackson, 1:2500) or HRP Conjugated goat anti mouse IgG (Jackson, 1:2500) then incubated 40 min at 4 °C. The plate was washed 3 times with 100 μl/well of the blocking buffer, 3 times with 100 μl/well washing buffer. 30 μl/well of BM Chemiluminescence ELISA substrate (Roche, 1:50) was added to the plate, and the luminosity was immediately measured using the Ensight Plate Reader.

### Flow cytometry

MRC-5 cells (0.4×10^6^ cells/well) were seeded in 6-well plates and after overnight growth transduced with Ad26 vectors encoding SARS-CoV-2 S transgenes at 5000 vp/cell for 48 hrs. Harvested cells were washed with PBS and stained with LIVE/DEAD^™^ Fixable Violet Dead Cell Stain Kit (Invitrogen). For SARS-CoV-2 surface staining, cells were washed twice with PBS and then incubated with ACE2-Fc (1 μg/ml), convalescent serum (1:400), and mAbs S309, SAD-S35, CR3022, CR3015 and CR3046 (1 μg/ml) for 30 min in FACS buffer (PBS with 0.5% BSA). Cells were washed twice with FACS buffer and stained with goat anti-Human IgG Alexa Fluor 647 (Invitrogen) or goat anti-Mouse IgG Alexa Fluor 647 (Invitrogen) secondary antibody for 30 min in FACS buffer. Cells were washed twice with FACS buffer and fixed with 1x BD CellFIX (BD Biosciences) for 15 min. Cells were washed once with FACS buffer and resuspended in FACS buffer before analysis on a FACS Canto instrument (BD Biosciences). Data were analyzed with FlowJo^™^ Software (Becton, Dickinson and Company) and plotted as median fluorescence intensity of the MRC-5 single, live cell population.

### BioLayer Interferometry (BLI)

Expi293F cells were transiently transfected using ExpiFectamine (Life Technologies) according to the manufacturer’s instructions and cultured for 3 days at 37°C and 10% CO2. The culture supernatant was harvested and spun for 5 minutes at 300 g to remove cells and cellular debris. The spun supernatant was subsequently sterile filtered using a 0.22 μm vacuum filter and used as the analyte in the experimentA solution of ACE2-Fc at a concentration of 10 μg/mL was used to immobilize the ligand on anti-hIgG (AHC) sensors (FortéBio cat#18-5060) in 1x kinetics buffer (FortéBio cat#18-1092) in 96-well black flat bottom polypropylene microplates (FortéBio cat#3694). The experiment was performed on an Octet HTX instrument (Pall-FortéBio) at 30 °C with a shaking speed of 1,000 rpm. Activation was 60 s, immobilization of antibodies 600 s, followed by washing for 300 s and then binding the S proteins for 1200 s. Data analysis was performed using the FortéBio Data Analysis 8.1 software (FortéBio). Binding levels were plotted as nm shifts at 1200s after S protein binding.

### Differential scanning fluorometry (DSF)

0.2 mg of purified protein in 50 µl PBS pH 7.4 (Gibco) was mixed with 15 µl of 20 times diluted SYPRO orange fluorescent dye (5000 x stock, Invitrogen S6650) in a 96-well optical qPCR plate. A negative control sample containing the dye only was used for reference subtraction. The measurement was performed in a qPCR instrument (Applied Biosystems ViiA 7) using a temperature ramp from 25–95°C with a rate of 0.015°C per second. Data were retrieved continuously. The negative first derivative was plotted as a function of temperature. The melting temperature corresponds to the lowest point in the curve.

### Mass-spectrometry

Liquid chromatography-mass spectrometry (LC-MS/MS) was used to determine the N-terminus on either purified soluble protein or on a pull-down of the full-length S protein from cell membranes using either Mab CR3022 or ACE2-Fc. The purified soluble proteins were subjected to direct digest, whereas the membrane-bound spike protein samples were either subjected to direct digest or in gel digestion. Data processing for the different proteins was performed using Biopharma finder 3.1 (Thermo Scientific). Each data file was compared to its corresponding amino acid sequence. Peptides were filtered by mass accuracy, confidence and structural resolution and reported. In order to pick up low abundant peptides, the thresholds for the MS noise level and the S/N were lowered 100-fold in comparison to the normal processing method.

### Cell-cell fusion assay

Cell-cell fusion assays were performed to ascertain the relative fusogenicity of the different S protein variants. For fluorescent read-out, full-length wildtype SARS-CoV-2 S protein and variants thereof, human ACE2, human TMPRSS2 and GFP were co-expressed from pcDNA2004 plasmids in HEK293 cells using Trans-IT transfection reagent according to the manufacturer’s instructions. Transfections were performed on 70% confluent cell monolayers in 6-well plates. Transfected cells were incubated at 37°C and 10% CO_2_ for 24 hrs before imaging on an EVOS cell imaging system (Thermofisher). Overlays between brightfield and GFP channels were made in ImageJ.

The fluorescent fusion assay was adapted to allow quantitative measurement of cell-cell fusion by leveraging the NanoBiT complementation system (Promega). Donor HEK293 cells were transfected with full-length S and the 11S subunit in 96-well white flat bottom TC-treated microtest assay plates. Acceptor HEK293 cells were transfected in 6-well plates (Corning) with ACE2, TMPRSS2 and the PEP86 subunit, or just the PEP86 subunit (‘No spike’) as negative control. 18hr after transfection, the acceptor cells were released by 0.1% trypsin/EDTA and added to the donor cells at a 1:1 ratio for 4 hr. Luciferase complementation was measured by incubating with Nano-Glo® Live Cell Reagent for 3 min, followed by read-out on an Ensight plate reader (PerkinElmer).

### Ad26 Vectors

Replication-incompetent, E1/E3-deleted Ad26-vectors were engineered using the AdVac system (Abbink et al., 2007), here using a single plasmid technology containing the Ad26 vector genome including a transgene expression cassette. The codon optimized SARS-COV-2 Spike genes (*as described in vaccine design*) were inserted into the E1-position of the Ad26 vector genomes under transcriptional control of the human cytomegalovirus promoter and the SV-40 polyadenylation sequence. Rescue and manufacturing of the Ad26 vectors was performed in the complementing cell line PER.C6 TetR (Wunderlich et al., 2018; Zahn et al., 2012). Virus particle (vp) titers in the Ad26 vector preparations were quantified by measurement of optical density at 260 nm (Maizel et al., 1968) and infectivity was assessed by quantitative potency assay (QPA)(Wang et al., 2005), using PER.C6 TetR cells. PCR including subsequent sequencing of PCR products has confirmed the identity and integrity of the SARS-COV-2 Spike genes. Ad26 vector-mediated expression of SARS-COV-2 Spike genes was confirmed by western blot analysis of cell-culture lysates from infected MRC-5 cells (Fig. 1G). Bioburden levels and endotoxin levels met the preset release criteria for animal experiments.

### SDS-PAGE and Western Blotting

24 well plates were seeded with MRC-5 cells (1.25×10^5^ cells/well), and after overnight growth transduced with Ad26 vectors encoding SARS-CoV-2 S transgenes. Cell lysates were harvested 48 hrs post transduction and, after heating for 5 min at 85°C, samples were loaded under non-reduced conditions on a precasted 3-8% Tris-Acetate SDS-PAGE gel (Invitrogen). Proteins were transferred to a nitrocellulose membrane using an iBlot2 dry blotting system (Invitrogen), and membrane blocking was performed overnight at 4°C in Tris-buffered saline (TBS) containing 0.2% Tween 20 (V/V) (TBST) and 5% (W/V) Blotting-Grade Blocker (Bio-Rad). Following overnight blocking, the membrane was incubated for 1 hr with 2.8 µg/ml CR3046. in TBST-5% Blocker. CR3046 is a human monoclonal antibody directed against SARS-CoV-1 Spike and binds to the Spike S2 domain and also cross-reacts with SARS-CoV-2 Spike S2 (unpublished data). After incubation, the membrane was washed three times with TBST for 5 min and subsequently incubated for 1 hr with 1:10,000 IRDye 800CW-conjugated goat-anti-human secondary antibody (Li-COR) in TBST-5% Blocker. Finally, the PVDF membrane was washed three times with TBST for 5 min, and after drying developed using an ODYSSEY® CLx Infrared Imaging System (Li-COR).

### Virus Neutralization Assay

Neutralization assays against live SARS-CoV-2 were performed using the microneutralization assay previously described by Algaissi and Hashem (Algaissi and Hashem, 2020). Vero E6 cells [CRL-1580, American Type Culture Collection (ATCC)] were grown in Eagle’s minimal essential medium (EMEM; Lonza) supplemented with 8% fetal calf serum (FCS; Bodinco BV), 1% penicillin-streptomycin (Sigma-Aldrich, P4458) and 2 mM L-glutamine (PAA). Cells were maintained at 37°C in a humidified atmosphere containing 5% CO_2_. Clinical isolate SARS-CoV-2/human/NLD/Leiden-0001/2020 (Leiden L-0001) was isolated from a nasopharyngeal sample and its characterization will be described elsewhere (manuscript in preparation). The NGS-derived sequence of this virus isolate is available under GenBank accession number MT705205. Isolate Leiden-0001 was propagated and titrated in Vero E6 cells using the tissue culture infective dose 50 (TCID_50_) endpoint dilution method and the TCID50 was calculated by the Spearman-Kärber algorithm as described in Hierholzer & Killington (1996) (Hierholzer and Killington, 1996). All work with live SARS-CoV-2 was performed in a biosafety level 3 facility at Leiden University Medical Center.

Vero-E6 cells were seeded at 12,000 cells/well in 96-well tissue culture plates 1 day prior to infection. Heat-inactivated (30 min at 56°C) serum samples were analyzed in duplicate. The panel of sera were two-fold serially diluted in duplicate, with an initial dilution of 1:10 and a final dilution of 1:1280 in 60 μL EMEM medium supplemented with penicillin, streptomycin, 2 mM L-glutamine and 2% FCS. Diluted sera were mixed with equal volumes of 120 TCID_50_/60 µL Leiden -0001 virus and incubated for 1 h at 37 °C. The virus-serum mixtures were then added onto Vero-E6 cell monolayers and incubated at 37 °C in a humidified atmosphere with 5 % CO_2_. Cells either unexposed to the virus or mixed with 120 TCID_50_/60 µL SARS-CoV-2 were used as negative (uninfected) and positive (infected) controls, respectively. At 3 days post-infection, cells were fixed and inactivated with 40 µL 37% formaldehyde/PBS solution/well overnight at 4 °C. The fixative was removed from cells and the clusters were stained with 50 µL/well crystal violet solution, incubated for 10 minutes and rinsed with water. Dried plates were evaluated for viral cytopathic effect. Neutralization titer was calculated by dividing the number of positive wells with complete inhibition of the virus-induced cytopathogenic effect, by the number of replicates, and adding 2.5 to stabilize the calculated ratio. The neutralizing antibody titer was defined as the log2 reciprocal of this value. A SARS-CoV-2 back-titration was included with each assay run to confirm that the dose of the used inoculum was within the acceptable range of 30 to 300 TCID_50_.

### Pseudotyped virus neutralization assay

MLV pseudotyped with SARS-COV-2 S protein was produced as previously described (Millet and Whittaker, 2016) with some minor changes. In short, Platinum-GP cells (cell Biolabs, Inc) were transfected with a plasmid encoding the codon optimized SARS-COV-2 Spike gene from strain Wuhan-Hu-1 (GenBank: MN908947) carrying a 19-aa cytoplasmic tail truncation, a GAG-Pol packaging vector and an MLV vector encoding a luciferase reporter gene using Lipofectamine 3000 transfection reagent (Life Technologies) according to manufacturer’s protocol. Cells were incubated overnight at 37°C 10% CO_2_ with OPTIMEM transfection medium. Next day, medium was replaced by OPTIMEM supplemented with 5% FBS and 1% PenStrep and incubated for 48 hrs prior to harvest. The harvested pseudotyped MLV particles were stored at -80°C prior to use. In the neutralization assay, soluble ACE2-Fc and mAbs CR3015, CR3022, CR3046 and SAD-S35 (ACROBiosystems) were two-fold serial diluted (n=3) in DMEM without phenol red supplemented with 1% FBS and 1% PenStrep and incubated with an equal volume of pseudotyped MLV. After 30 min incubation, the mixture was inoculated onto susceptible Vero E6 cells in MW96 well plates. Luciferase activity was measured after 40 h of incubation by addition of an equal volume of substrate NeoLite (Perkin Elmer) followed by luminescence read out on the EnSight Multimode Plate Reader (Perkin Elmer). The percentage of infectivity was calculated as ratio of luciferase readout in the presence of mAbs normalized to luciferase readout in the absence of mAb.

### Direct coat ELISAs

IgG binding to SARS-CoV-2 Spike antigen was measured by ELISA with the full-length in house produced Spike protein COR200099. COR200099 is an in-house produced SARS-CoV-2 Spike protein, stabilized by two point mutations in the S1/S2 junction that knocks out the furin cleavage site, and by two newly introduced prolines in the hinge region in S2. In addition, the transmembrane and cytoplasmic regions have been replaced by a foldon domain for trimerization mutations, allowing the protein to be produced as soluble protein (see S.dTM.PP Fig. 3C). Briefly, 96-wells Perkin Elmer white ½ area plates were O/N coated with COR200099 protein. Next day plates were washed, blocked for 1 hr and subsequently incubated for 1 hr with 3-fold serially diluted serum samples in block buffer in duplicate. After washing, plates were incubated for 1 hr with goat-anti-mouse IgG-HRP in block buffer, washed again and developed using ECL substrate. Luminescence readout was performed using a BioTek Synergy Neo instrument.

For IgG1 and IgG2a ELISAs, a similar protocol was followed as described above, but respectively using Goat anti-Mouse IgG1-HRP and Goat anti-Mouse IgG2a-HRP as secondary antibodies.

### ELISpot

IFN-γ ELISpot was performed on splenocytes of mice isolated after sacrifice using mouse IFN-γ ELISpot-plus kit (Mabtech). Splenocytes were obtained by disaggregation of spleens with the gentleMACS dissociator. IFN-γ ELISpot assay was performed by stimulating splenocytes from individual mice for 18 h with two different peptide pools (pool 1; peptides 1-156, and pool 2; peptides 157-313) consisting of in total 313 15-mers peptides overlapping by 11 amino acids together covering the full-length Spike protein at a final concentration of 1 μg/peptide/mL. Results shared in this manuscript are the sum of stimulation with peptide pool 1 and pool 2. PMA/ionomycin stimulation was used as a positive control (data not shown); medium was used as negative control and used to calculate the lower limit of detection. Stimulation was done overnight in duplicate wells containing 0.5 - 2.5 ⨯ 10^5^ cells per well.

### Multiplex ELISA

The Th1/Th2 multiplex ELISA assay was performed on splenocytes obtained after sacrifice. Splenocytes were stimulated by 48 h culturing in the presence of two Spike 15-mer peptide pools (pool 1 and pool 2). Resulting supernatants were diluted 4-fold and measured for the presence of Th1 (IFN-γ) and Th2 cytokines (IL-10, IL-4 and IL-5) using a 10-plex multiplex ELISA pro-inflammatory panel (mouse) kit (Meso Scale Discovery, V-PLEX Proinflammatory Panel 1 Mouse Kit cat# K15048D). Results shared in this manuscript are the sum of stimulation with peptide pool 1 and pool 2. Ratios of Th1/Th2 cytokines (IFN-γ/IL-10, IFN-γ/IL-4 and IFN-γ/IL-5) were calculated on basis of cytokine measurements in supernatants by dividing the Th1 cytokine (IFN-γ) with the respective Th2 cytokines. Multiplex ELISA measurements were done on supernatant diluted either 2-fold or 4-fold.

### Animals

Female BALB/c or C57BL6 mice (specific pathogen-free), aged 8-12 weeks at the start of the study were purchased from Charles River laboratories (Sulzfeld, Germany). Mice were immunized via the intramuscular (i.m.) route with 100μl vaccine (50μl per hind leg) under isoflurane anesthesia. Intermediate blood samples were collected via submandibular bleeding route. At the end of the experiment, under anesthesia, animals were exsanguinated by heart puncture and sacrificed by cervical dislocation. Blood was processed for serum isolation and spleens were collected. Spleens were processed into single cell suspensions for cellular assays (when applicable).

### Statistical analysis

Statistical differences between immunization regimens were evaluated for S-specific binding antibodies as measured by ELISA, NAb titers as measured by VNA, IFN-γ producing cells as measured by ELISpot and cytokine production by MSD and ICS assays. Comparisons between Ad26.S, Ad26.tPA.S, Ad26.tPA.SS, Ad26.tPA.WT.S, Ad26.S.PP, adjuvanted S protein, tPA.S, tPA.S.PP, S and S.PP groups were made using the exact Wilcoxon rank-sum test, Cochran-Mantel-Haenszel test, Mann-Whitney U test, t-test from ANOVA, or z-test from Tobit ANOVA. Results were corrected for multiple comparisons by Bonferroni correction; 3-fold Bonferroni correction Figures 5 A and B, 2-fold Bonferroni correction Figures 5 C and D.

Statistical analyses were performed using SAS version 9.4 (SAS Institute Inc. Cary, NC, US) and R version 3.6.1 (2019-07-05). Statistical tests were conducted two-sided at an overall significance level of α = 0.05.

## References

Abbink, P., Lemckert, A.A., Ewald, B.A., Lynch, D.M., Denholtz, M., Smits, S., Holterman, L., Damen, I., Vogels, R., Thorner, A.R., et al. (2007). Comparative seroprevalence and immunogenicity of six rare serotype recombinant adenovirus vaccine vectors from subgroups B and D. J Virol 81, 4654–4663.

Algaissi, A., and Hashem, A.M. (2020). Evaluation of MERS-CoV Neutralizing Antibodies in Sera Using Live Virus Microneutralization Assay. Methods Mol Biol 2099, 107–116.

Alharbi, N.K., Padron-Regalado, E., Thompson, C.P., Kupke, A., Wells, D., Sloan, M.A., Grehan, K., Temperton, N., Lambe, T., Warimwe, G., et al. (2017). ChAdOx1 and MVA based vaccine candidates against MERS-CoV elicit neutralising antibodies and cellular immune responses in mice. Vaccine 35, 3780–3788.

Anywaine, Z., Whitworth, H., Kaleebu, P., Praygod, G., Shukarev, G., Manno, D., Kapiga, S., Grosskurth, H., Kalluvya, S., Bockstal, V., et al. (2019). Safety and Immunogenicity of a 2-Dose Heterologous Vaccination Regimen With Ad26.ZEBOV and MVA-BN-Filo Ebola Vaccines: 12-Month Data From a Phase 1 Randomized Clinical Trial in Uganda and Tanzania. J Infect Dis 220, 46–56.

Baden, L.R., Karita, E., Mutua, G., Bekker, L.G., Gray, G., Page-Shipp, L., Walsh, S.R., Nyombayire, J., Anzala, O., Roux, S., et al. (2016). Assessment of the Safety and Immunogenicity of 2 Novel Vaccine Platforms for HIV-1 Prevention: A Randomized Trial. Ann Intern Med 164, 313–322.

Baden, L.R., Walsh, S.R., Seaman, M.S., Tucker, R.P., Krause, K.H., Patel, A., Johnson, J.A., Kleinjan, J., Yanosick, K.E., Perry, J., et al. (2013). First-in-human evaluation of the safety and immunogenicity of a recombinant adenovirus serotype 26 HIV-1 Env vaccine (IPCAVD 001). J Infect Dis 207, 240–247.

Barouch, D.H., Liu, J., Peter, L., Abbink, P., Iampietro, M.J., Cheung, A., Alter, G., Chung, A., Dugast, A.S., Frahm, N., et al. (2013). Characterization of humoral and cellular immune responses elicited by a recombinant adenovirus serotype 26 HIV-1 Env vaccine in healthy adults (IPCAVD 001). J Infect Dis 207, 248–256.

Barouch, D.H., Tomaka, F.L., Wegmann, F., Stieh, D.J., Alter, G., Robb, M.L., Michael, N.L., Peter, L., Nkolola, J.P., Borducchi, E.N., et al. (2018). Evaluation of a mosaic HIV-1 vaccine in a multicentre, randomised, double-blind, placebo-controlled, phase 1/2a clinical trial (APPROACH) and in rhesus monkeys (NHP 13-19). Lancet 392, 232–243.

Battles, M.B., Mas, V., Olmedillas, E., Cano, O., Vazquez, M., Rodriguez, L., Melero, J.A., and McLellan, J.S. (2017). Structure and immunogenicity of pre-fusion-stabilized human metapneumovirus F glycoprotein. Nat Commun 8, 1528.

Chandrashekar, A., Liu, J., Martinot, A.J., McMahan, K., Mercado, N.B., Peter, L., Tostanoski, L.H., Yu, J., Maliga, Z., Nekorchuk, M., et al. (2020). SARS-CoV-2 infection protects against rechallenge in rhesus macaques. Science.

Corbett, K.S., Edwards, D., Leist, S.R., Abiona, O.M., Boyoglu-Barnum, S., Gillespie, R.A., Himansu, S., Schafer, A., Ziwawo, C.T., DiPiazza, A.T., et al. (2020). SARS-CoV-2 mRNA Vaccine Development Enabled by Prototype Pathogen Preparedness. bioRxiv.

Coronaviridae Study Group of the International Committee on Taxonomy of, V. (2020). The species Severe acute respiratory syndrome-related coronavirus: classifying 2019-nCoV and naming it SARS-CoV-2. Nat Microbiol 5, 536–544.

Gilbert, S.C., Hill, A.V.S., and Moris, S.J. (2017). COMPOSITIONS AND METHODS FOR INDUCING AN IMMUNE RESPONSE. patent WO 2018/215766 Al

Hastie, K.M., Zandonatti, M.A., Kleinfelter, L.M., Heinrich, M.L., Rowland, M.M., Chandran, K., Branco, L.M., Robinson, J.E., Garry, R.F., and Saphire, E.O. (2017). Structural basis for antibody-mediated neutralization of Lassa virus. Science 356, 923–928.

Hierholzer, J.C., and Killington, R.A. (1996). Virus isolation and quantitation. Virology Methods Manual 1st edition, 25–46.

Honda-Okubo, Y., Barnard, D., Ong, C.H., Peng, B.H., Tseng, C.T., and Petrovsky, N. (2015). Severe acute respiratory syndrome-associated coronavirus vaccines formulated with delta inulin adjuvants provide enhanced protection while ameliorating lung eosinophilic immunopathology. J Virol 89, 2995–3007.

Iwata-Yoshikawa, N., Uda, A., Suzuki, T., Tsunetsugu-Yokota, Y., Sato, Y., Morikawa, S., Tashiro, M., Sata, T., Hasegawa, H., and Nagata, N. (2014). Effects of Toll-like receptor stimulation on eosinophilic infiltration in lungs of BALB/c mice immunized with UV-inactivated severe acute respiratory syndrome-related coronavirus vaccine. J Virol 88, 8597–8614.

Kirchdoerfer, R.N., Wang, N., Pallesen, J., Wrapp, D., Turner, H.L., Cottrell, C.A., Corbett, K.S., Graham, B.S., McLellan, J.S., and Ward, A.B. (2018). Stabilized coronavirus spikes are resistant to conformational changes induced by receptor recognition or proteolysis. Sci Rep 8, 15701.

Krarup, A., Truan, D., Furmanova-Hollenstein, P., Bogaert, L., Bouchier, P., Bisschop, I.J.M., Widjojoatmodjo, M.N., Zahn, R., Schuitemaker, H., McLellan, J.S., et al. (2015). A highly stable prefusion RSV F vaccine derived from structural analysis of the fusion mechanism. Nat Commun 6, 8143.

Lindblad, E.B. (2004). Aluminium compounds for use in vaccines. Immunol Cell Biol 82, 497–505.

Liu, J., Ewald, B.A., Lynch, D.M., Denholtz, M., Abbink, P., Lemckert, A.A., Carville, A., Mansfield, K.G., Havenga, M.J., Goudsmit, J., et al. (2008). Magnitude and phenotype of cellular immune responses elicited by recombinant adenovirus vectors and heterologous prime-boost regimens in rhesus monkeys. J Virol 82, 4844–4852.

Liu, P., Wysocki, J., Souma, T., Ye, M., Ramirez, V., Zhou, B., Wilsbacher, L.D., Quaggin, S.E., Batlle, D., and Jin, J. (2018). Novel ACE2-Fc chimeric fusion provides long-lasting hypertension control and organ protection in mouse models of systemic renin angiotensin system activation. Kidney Int 94, 114–125.

Maizel, J.V., Jr., White, D.O., and Scharff, M.D. (1968). The polypeptides of adenovirus. I. Evidence for multiple protein components in the virion and a comparison of types 2, 7A, and 12. Virology 36, 115–125.

Martin, J.E., Louder, M.K., Holman, L.A., Gordon, I.J., Enama, M.E., Larkin, B.D., Andrews, C.A., Vogel, L., Koup, R.A., Roederer, M., et al. (2008). A SARS DNA vaccine induces neutralizing antibody and cellular immune responses in healthy adults in a Phase I clinical trial. Vaccine 26, 6338–6343.

Millet, J.K., and Whittaker, G.R. (2016). Murine Leukemia Virus (MLV)-based Coronavirus Spike-pseudotyped Particle Production and Infection. Bio Protoc 6.

Milligan, I.D., Gibani, M.M., Sewell, R., Clutterbuck, E.A., Campbell, D., Plested, E., Nuthall, E., Voysey, M., Silva-Reyes, L., McElrath, M.J., et al. (2016). Safety and Immunogenicity of Novel Adenovirus Type 26- and Modified Vaccinia Ankara-Vectored Ebola Vaccines: A Randomized Clinical Trial. JAMA 315, 1610–1623.

Mutua, G., Anzala, O., Luhn, K., Robinson, C., Bockstal, V., Anumendem, D., and Douoguih, M. (2019). Safety and Immunogenicity of a 2-Dose Heterologous Vaccine Regimen With Ad26.ZEBOV and MVA-BN-Filo Ebola Vaccines: 12-Month Data From a Phase 1 Randomized Clinical Trial in Nairobi, Kenya. J Infect Dis 220, 57–67.

Pallesen, J., Wang, N., Corbett, K.S., Wrapp, D., Kirchdoerfer, R.N., Turner, H.L., Cottrell, C.A., Becker, M.M., Wang, L., Shi, W., et al. (2017). Immunogenicity and structures of a rationally designed prefusion MERS-CoV spike antigen. Proc Natl Acad Sci U S A 114, E7348–E7357.

Pinto, D., Park, Y.J., Beltramello, M., Walls, A.C., Tortorici, M.A., Bianchi, S., Jaconi, S., Culap, K., Zatta, F., De Marco, A., et al. (2020). Cross-neutralization of SARS-CoV-2 by a human monoclonal SARS-CoV antibody. Nature 583, 290–295.

Radosevic, K., Rodriguez, A., Lemckert, A.A., van der Meer, M., Gillissen, G., Warnar, C., von Eyben, R., Pau, M.G., and Goudsmit, J. (2010). The Th1 immune response to Plasmodium falciparum circumsporozoite protein is boosted by adenovirus vectors 35 and 26 with a homologous insert. Clin Vaccine Immunol 17, 1687–1694.

Rutten, L., Gilman, M.S.A., Blokland, S., Juraszek, J., McLellan, J.S., and Langedijk, J.P.M. (2020). Structure-Based Design of Prefusion-Stabilized Filovirus Glycoprotein Trimers. Cell Rep 30, 4540–4550 e4543.

Rutten, L., Lai, Y.T., Blokland, S., Truan, D., Bisschop, I.J.M., Strokappe, N.M., Koornneef, A., van Manen, D., Chuang, G.Y., Farney, S.K., et al. (2018). A Universal Approach to Optimize the Folding and Stability of Prefusion-Closed HIV-1 Envelope Trimers. Cell Rep 23, 584–595.

Sanders, R.W., Vesanen, M., Schuelke, N., Master, A., Schiffner, L., Kalyanaraman, R., Paluch, M., Berkhout, B., Maddon, P.J., Olson, W.C., et al. (2002). Stabilization of the soluble, cleaved, trimeric form of the envelope glycoprotein complex of human immunodeficiency virus type 1. J Virol 76, 8875–8889.

Shukarev, G., Callendret, B., Luhn, K., Douoguih, M., and consortium, E. (2017). A two-dose heterologous prime-boost vaccine regimen eliciting sustained immune responses to Ebola Zaire could support a preventive strategy for future outbreaks. Hum Vaccin Immunother 13, 266–270.

Smith, T.R.F., Patel, A., Ramos, S., Elwood, D., Zhu, X., Yan, J., Gary, E.N., Walker, S.N., Schultheis, K., Purwar, M., et al. (2020). Immunogenicity of a DNA vaccine candidate for COVID-19. Nat Commun 11, 2601.

Sternberg, A., and Naujokat, C. (2020). Structural features of coronavirus SARS-CoV-2 spike protein: Targets for vaccination. Life Sci 257, 118056.

Tortorici, M.A., and Veesler, D. (2019). Structural insights into coronavirus entry. Adv Virus Res 105, 93–116.

Tseng, C.T., Sbrana, E., Iwata-Yoshikawa, N., Newman, P.C., Garron, T., Atmar, R.L., Peters, C.J., and Couch, R.B. (2012). Immunization with SARS coronavirus vaccines leads to pulmonary immunopathology on challenge with the SARS virus. PLoS One 7, e35421.

van den Brink, E.N., Ter Meulen, J., Cox, F., Jongeneelen, M.A., Thijsse, A., Throsby, M., Marissen, W.E., Rood, P.M., Bakker, A.B., Gelderblom, H.R., et al. (2005). Molecular and biological characterization of human monoclonal antibodies binding to the spike and nucleocapsid proteins of severe acute respiratory syndrome coronavirus. J Virol 79, 1635–1644.

van der Fits, L., Bolder, R., Heemskerk-van der Meer, M., Drijver, J., van Polanen, Y., Serroyen, J., Langedijk, J.P.M., Schuitemaker, H., Saeland, E., and Zahn, R. (2020). Adenovector 26 encoded prefusion conformation stabilized RSV-F protein induces long-lasting Th1-biased immunity in neonatal mice. NPJ Vaccines 5, 49.

van Doremalen, N., Lambe, T., Spencer, A., Belij-Rammerstorfer, S., Purushotham, J.N., Port, J.R., Avanzato, V., Bushmaker, T., Flaxman, A., Ulaszewska, M., et al. (2020). ChAdOx1 nCoV-19 vaccination prevents SARS-CoV-2 pneumonia in rhesus macaques. bioRxiv.

Walls, A.C., Park, Y.J., Tortorici, M.A., Wall, A., McGuire, A.T., and Veesler, D. (2020). Structure, Function, and Antigenicity of the SARS-CoV-2 Spike Glycoprotein. Cell 181, 281–292 e286.

Wang, F., Puddy, A.C., Mathis, B.C., Montalvo, A.G., Louis, A.A., McMackin, J.L., Xu, J., Zhang, Y., Tan, C.Y., Schofield, T.L., et al. (2005). Using QPCR to assign infectious potencies to adenovirus based vaccines and vectors for gene therapy: toward a universal method for the facile quantitation of virus and vector potency. Vaccine 23, 4500–4508.

Winslow, R.L., Milligan, I.D., Voysey, M., Luhn, K., Shukarev, G., Douoguih, M., and Snape, M.D. (2017). Immune Responses to Novel Adenovirus Type 26 and Modified Vaccinia Virus Ankara-Vectored Ebola Vaccines at 1 Year. JAMA 317, 1075–1077.

Wrapp, D., Wang, N., Corbett, K.S., Goldsmith, J.A., Hsieh, C.L., Abiona, O., Graham, B.S., and McLellan, J.S. (2020). Cryo-EM structure of the 2019-nCoV spike in the prefusion conformation. Science 367, 1260–1263.

Wunderlich, K., Uil, T., Gilles, Vellinga J., Sanders, B. Petronella, and Van der Vlugt, R. (2018). POTENT AND SHORT PROMOTER FOR EXPRESSION OF HETEROLOGOUS GENES. patent WO/2018/146205.

Yu, J., Tostanoski, L.H., Peter, L., Mercado, N.B., McMahan, K., Mahrokhian, S.H., Nkolola, J.P., Liu, J., Li, Z., Chandrashekar, A., et al. (2020). DNA vaccine protection against SARS-CoV-2 in rhesus macaques. Science.

Zahn, R., Gillisen, G., Roos, A., Koning, M., van der Helm, E., Spek, D., Weijtens, M., Grazia Pau, M., Radosevic, K., Weverling, G.J., et al. (2012). Ad35 and ad26 vaccine vectors induce potent and cross-reactive antibody and T-cell responses to multiple filovirus species. PLoS One 7, e44115.

