## Supplementary figures for "Ad26-vector based COVID-19 vaccine encoding a prefusion stabilized SARS-CoV-2 Spike immunogen induces potent humoral and cellular immune responses": Bos et al COVID19 vaccine design suppl fig.pdf

Supplementary Figure 1.

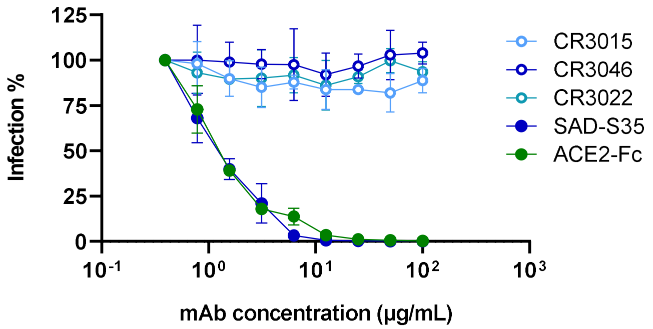


**Figure S1. Antibody-mediated neutralization of MLV particles pseudotyped with S protein of SARS-CoV-2.** Pseudotyped MLV particles pre-incubated with monoclonal antibodies or soluble receptor ACE2-Fc in a concentration range were used to transduce Vero E6 cells. The luminescence signal of the luciferase activity in transduced Vero E6 cells was measured 40 h post transduction to calculate the infection (%) of the Vero E6 cells relative to non-treated MLV particles.

Supplementary Figure 2.

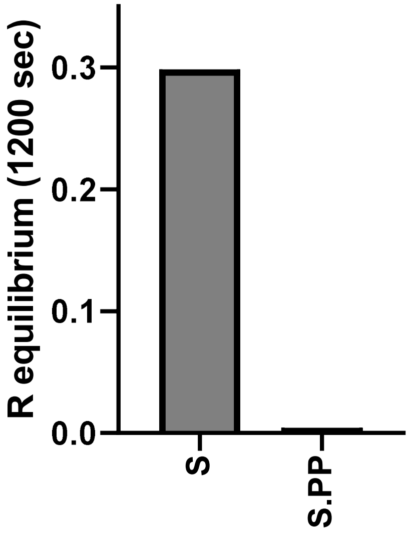


**Figure S2. Shedding of SARS-CoV-2 S1.** Biolayer interferometry to determine the binding of ACE2 to Expi293F cell culture supernatants expressing S and S.PP three days after transfection.

Supplementary Figure 3.

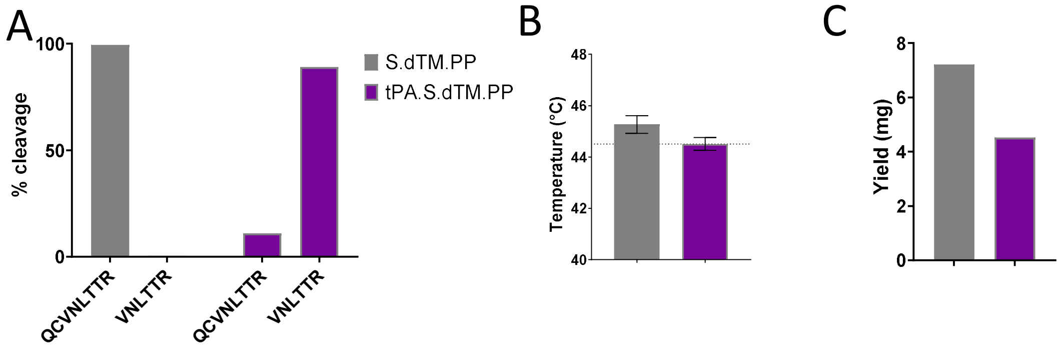


**Figure S3. Soluble protein characterization. A**) Percentage of SP cleavage at cleavage site 1 (CS1) resulting in peptide QCVNLTTR and at cleavage site 2 (CS2) resulting in peptide VNLTTR (see Fig 3A) as determined with mass-spectrometry. **B**) Melting temperature determined with DSF. The dotted horizontal line indicates the height of the $Tm_{50}$ of tPAs.S.dTM.PP. **C**) Protein yield per 900 ml culture for secreted S proteins.

Supplementary Figure 4.

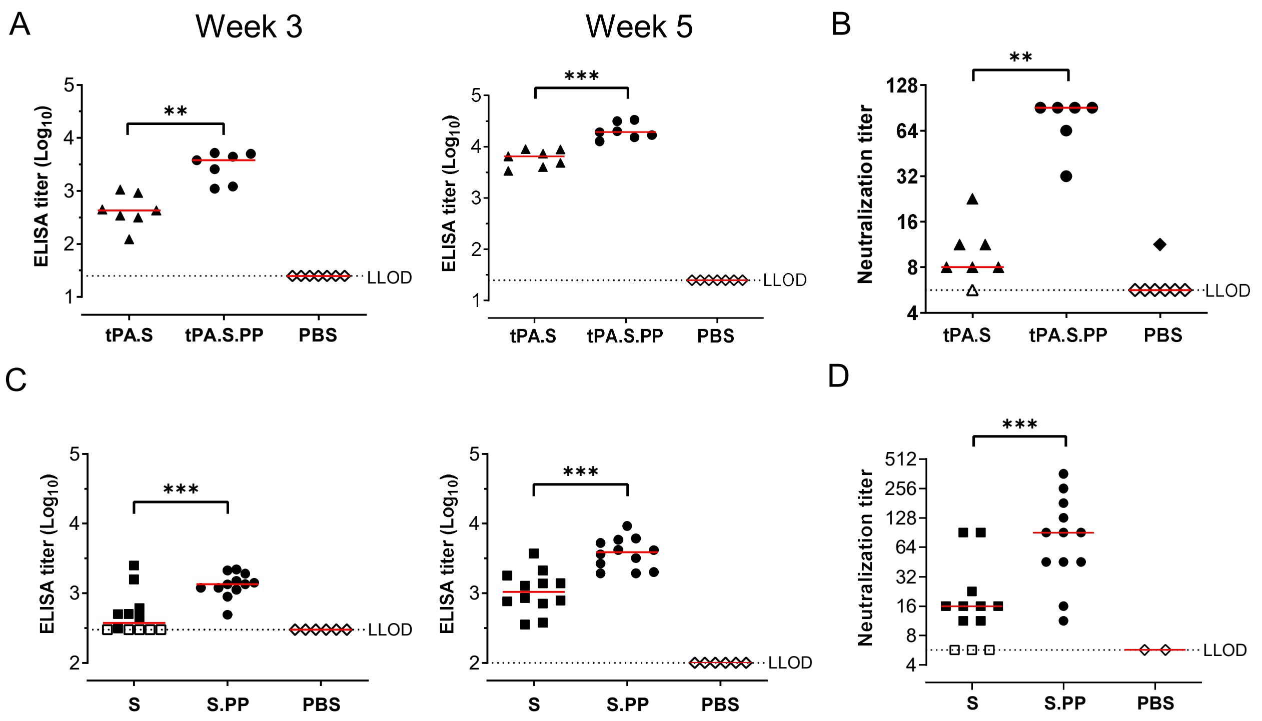


**Figure S4. S protein-specific antibody binding titers and SARS-CoV-2 NAbs induced by DNA immunization.** Naïve mice (BALB/c) were immunized twice, 3 weeks apart (Week 0 and Week 3), with 100 µg of DNA constructs. Serum samples were taken prior to the second immunization (Week 3), and 2 weeks after the second immunization (Week 5). Spike protein-specific binding and SARS-CoV-2 NAb titers were determined. In two separate experiments, mice were immunized with either A + B) tPA.S (N=7), tPA.S.PP (N=7), or PBS (N=7), or C + D) were immunized with either S (N=12), S.PP (N=12), or with PBS (N=6). Spike protein binding antibody titers were measured by ELISA. SARS-CoV-2 NAb titers were measured by wt VNA determining inhibition of the cytopathic effect (CPE) of virus isolate Leiden1 (L-0001) on Vero E6 cells. Mice immunized with PBS were taken along as two separate pools in fig S2D. The median response per group is indicated with a horizontal line. The dotted lines indicate LLODs. Animals with a response at or below the LLOD were put on LLOD and are shown as open symbols. Statistically significant differences indicated by asterisks; **: p<0.01, ***p<0.001.

Supplementary Figure 5.

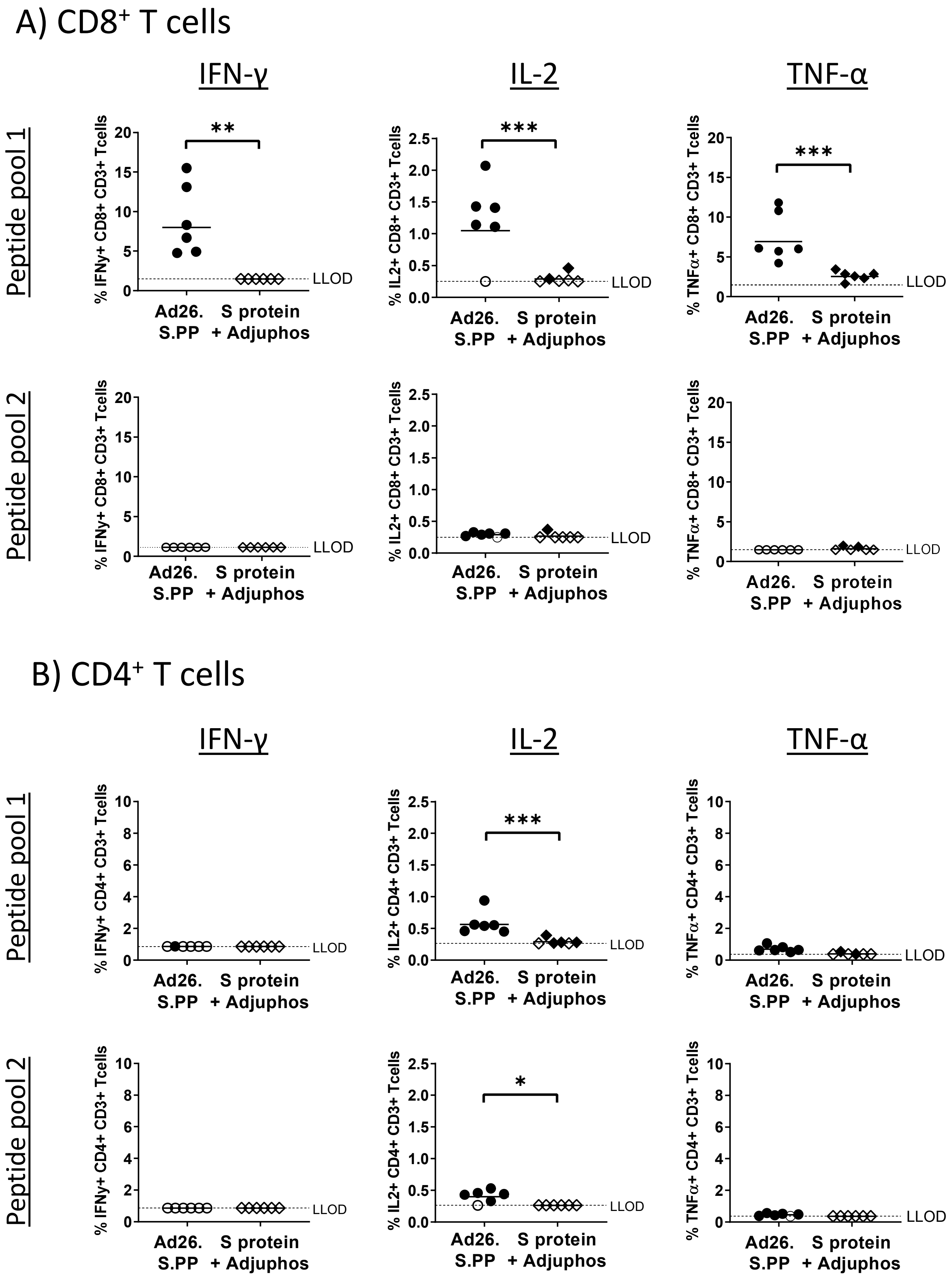

**Figure S5. Th1 associated cytokines induced by Ad26.S.PP compared with Adjuphos adjuvanted protein**. Naïve mice (BALB/c, N=6 per group) were immunized with either $10^{10}$ vp of Ad26.S.PP or 50 mcg of Spike protein adjuvanted with 100 µg Adjuphos (Adju-Phos®). Two weeks after immunization splenocytes were analyzed for intracellular cytokine expression of IFN-γ, IL-2 and TNF-α after stimulation with SARS-CoV-2 Spike protein peptides. A) CD8 positive T cells and B) CD4 positive T cells were stimulated with either S protein peptide pool 1 or peptide pool 2.

The median response per group is indicated with a horizontal line. The dotted lines indicate LLODs. Animals with a response at or below the LLOD were put on LLOD and are shown as open symbols. Statistically significant differences indicated by asterisks; *:p<0.05, **: p<0.01, ***p<0.001.
